# A retinal circuit generating a dynamic predictive code for orientated features

**DOI:** 10.1101/331504

**Authors:** Jamie Johnston, Sofie-Helene Seibel, Léa Simone Adele Darnet, Sabine Renninger, Michael Orger, Leon Lagnado

## Abstract

Sensory systems must reduce the transmission of redundant information to function efficiently. One strategy is to continuously adjust the sensitivity of neurons to suppress responses to common features of the input while enhancing responses to new ones. Here we image both the excitatory synaptic inputs and outputs of retinal ganglion cells to understand how such dynamic predictive coding is implemented in the analysis of spatial patterns. Synapses of bipolar cells become tuned to orientation through presynaptic inhibition generating lateral antagonism in the orientation domain. Individual ganglion cells receive excitatory synapses tuned to different orientations but feedforward inhibition generates a high-pass filter that only transmits the initial activation of these inputs, thereby removing redundancy. These results demonstrate how a dynamic predictive code can be implemented by circuit motifs common to many parts of the brain.

---

A general principle in understanding the design of sensory systems is the need to encode information efficiently, which in turn requires the removal of redundancies in the signal received from the outside world^1–4^. This principle helps us understand why the retina does not operate simply like a camera conveying a stream of intensity values for each pixel. Natural images contain a large amount of redundant information because pixels nearby in space and/or time tend to be correlated^5–7^. Rather than continuously transmitting the presence of an unchanging visual input, many retinal neurons preferentially signal deviations from the local image statistics, such as regions of contrast or the sudden movement of an object. Representing information by ignoring statistical regularities to highlight unusual components is termed predictive coding - an approach widely used in informatics to, for instance, compress images^8^. Examples of predictive coding have been identified throughout the brain, including the circuits involved in vision ^9–12^, hearing^13,14^ and touch^15^, the encoding of reward in the midbrain^16,17^ and space in the hippocampus and entorhinal cortex^18,19^. Although a number of models have been proposed for the neural implementation of predictive codes^9,20,21^, the circuit mechanisms have not been clearly identified.

Predictive coding can be a dynamic process, consistent with the animals need to adjust to sensory environments with different statistics^12,13^. For instance, the retina can adjust within seconds to changes in the variance in light intensity (contrast) by altering the strength of excitatory and inhibitory synapses^22,23^. A particularly striking example of adaptation occurs in response to changes in spatial correlations within the visual input, as are generated by edges of objects^24–26^. Neurons that encode orientation have been studied in detail in the visual cortex ^27–30^ but this information is already contained in the signals that retinal ganglion cells (RGCs) send back to the brain^31^. Further, some RGCs implement a dynamic predictive code (DPC) by rapidly altering their sensitivity to orientation in response to changes in the distribution of spatial correlations in the visual environment, becoming less sensitive to orientations that are common and more sensitive to orientations that are uncommon^9,32,33^.

The neural circuitry by which the visual system implements a DPC is not understood, either in the retina^23^ or in the cortex^20,26^. One model proposes the construction of a modifiable pattern detector that is fed by an array of excitatory subunits, each one tuned to a different stimulus pattern: if one of these detector subunits is driven strongly it fatigues and makes a smaller contribution to the activity of the output neuron which therefore becomes more sensitive to other, rarer, patterns^9,21,33,34^. This model is attractive but is not thought to operate in the retina because bipolar cells providing excitatory inputs to RGCs do not appear to be sensitive to orientation^9^. An alternative hypothesis of “network plasticity” has therefore been proposed in which the locus of the adaptive changes are the synapses that RGCs receive from the inhibitory amacrine cells, with these synapses obeying an anti-Hebbian plasticity rule that strengthens them when they are coactivated with the RGC^9^. In this scheme, a vertically orientated stimulus will strengthen inhibitory inputs above and below the RGC making it more sensitive to horizontal orientations.

To identify the circuit mechanisms generating dynamic predictive coding of spatial patterns we used an optical technique based on the fluorescent glutamate sensor iGluSnFR^35^ to compare the synaptic output from individual RGCs with the excitatory synaptic inputs that they receive from an array of bipolar cells. We probed these synapses using gratings of different orientations and found that many are strongly tuned to orientation through lateral antagonism in the orientation domain. Further, a subset of RGCs were driven by a mixture of excitatory synapses tuned to different orientations, providing the basic connectivity for a modifiable pattern detector. The nulling of a constant signal is then generated through at least two mechanisms: depression of excitatory synapses and fast feedforward inhibition. The basic features of this circuit are found in many other parts of the brain and may provide a general design for detecting change and removing redundancy.

## Imaging the input and output from individual retinal ganglion cells

The signals that RGCs deliver to the brain depend on the integration of a variety of synaptic inputs, often with mixed properties. Understanding how the activity of these inputs determine the output of the neuron has been difficult because individual synapses on a dendrite cannot be isolated using electrophysiology^36,37^. To overcome this problem we used *in vivo* imaging in larval zebrafish expressing iGluSnFR over the surface of subsets of RGCs (Fig. 1A). Visual stimuli generated “hotspots” of iGluSnFR fluorescence on dendrites (Fig. 1B) and these displayed many of the functional properties expected of glutamate release from the synapses of bipolar cells, such as adaptation to contrast^38^ (Fig. 2). The axons of these same RGCs could then be tracked to monitor the signal delivered in the optic tectum, also by the release of glutamate (Fig. 1A). iGluSnFR signals in the tectum were abolished when the cell body in the retina was ablated, demonstrating that they reflected the synaptic release of glutamate from the imaged neuron rather than glutamate spillover from neighbouring cells (Fig. 1C).

**Figure 1.**
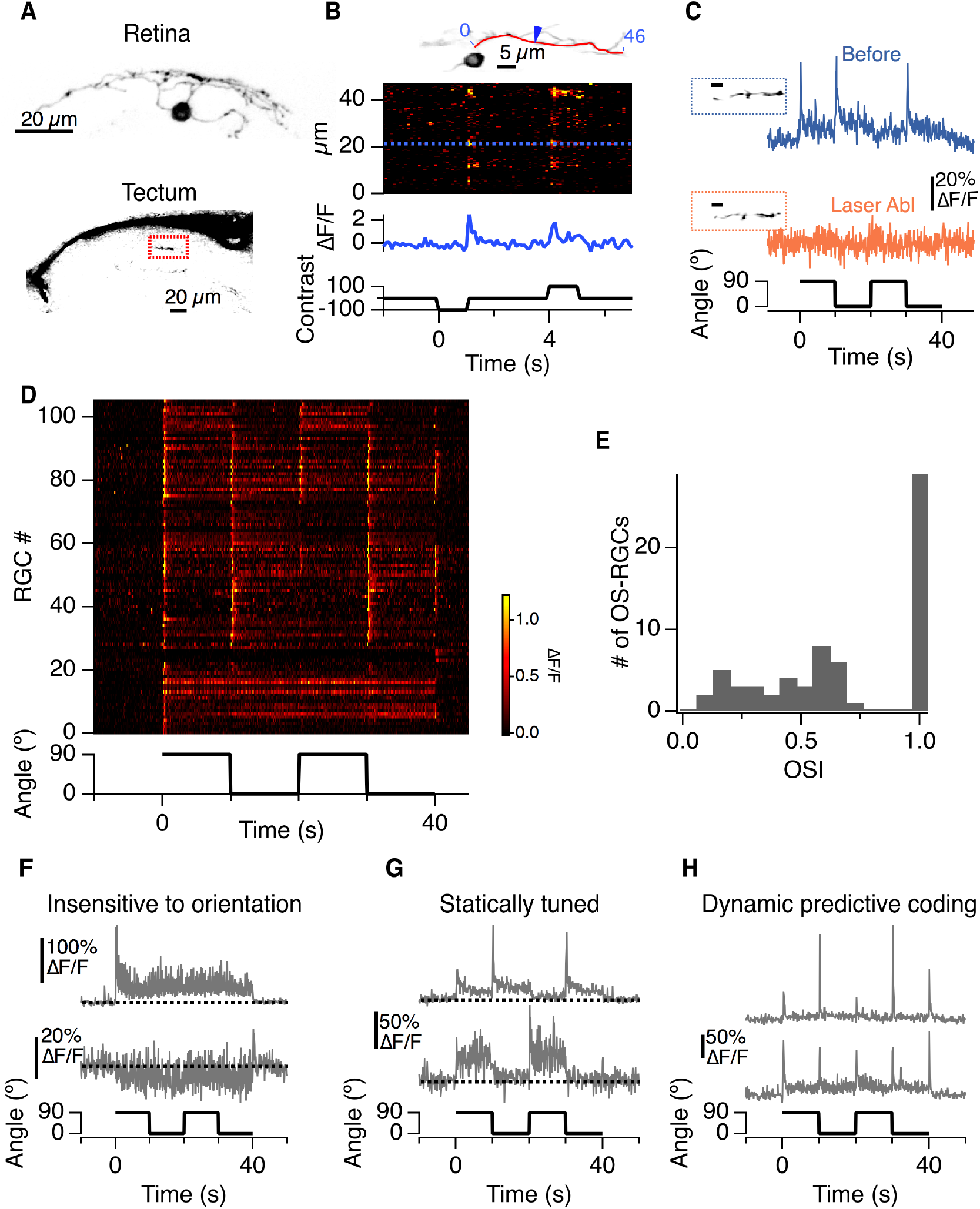
Imaging the input and output from individual retinal ganglion cells. **A**) Maximum intensity projection of an RGC labelled with iGluSNFR and imaged *in vivo* and its axon terminal in the optic tectum. **B**) *Top*: A single plane imaged through an RGC. *Middle:* A raster plot showing the time series of the iGluSnFR signal along a dendrite (redline) in response to changes in full field luminance. *Bottom*: Time course of the iGluSnFR signal at the point indicated by the blue arrow (corresponding to dashed blue line above). **C**) The iGluSnFR signal from an RGC axon terminal in response to a full-field grating reversing contrast at 5 Hz that switched orientation from 90° to 0° at 10 s intervals. The blue and orange traces show the responses before and after laser ablation of the RGC’s cell body. **D**) Raster plot of the responses of 106 retinal ganglion cell terminals to the contrast reversing grating. Neurons 0-27 only responded to changes in contrast. Neurons exhibiting DPC 72-96 responded with transient increases in glutamate at each change in stimulus orientation. **E**) Histogram of the OSI for the 66 RGCs in D showing significant orientation selectivity. **F**) Examples of output from RGCs that were insensitive to orientation. The lower example was inhibited by an increase in contrast. **G**) Examples of output from RGCs with static tuning to orientation. **H**). Examples of output from two RGCs displaying DPC.

**Figure 2.**
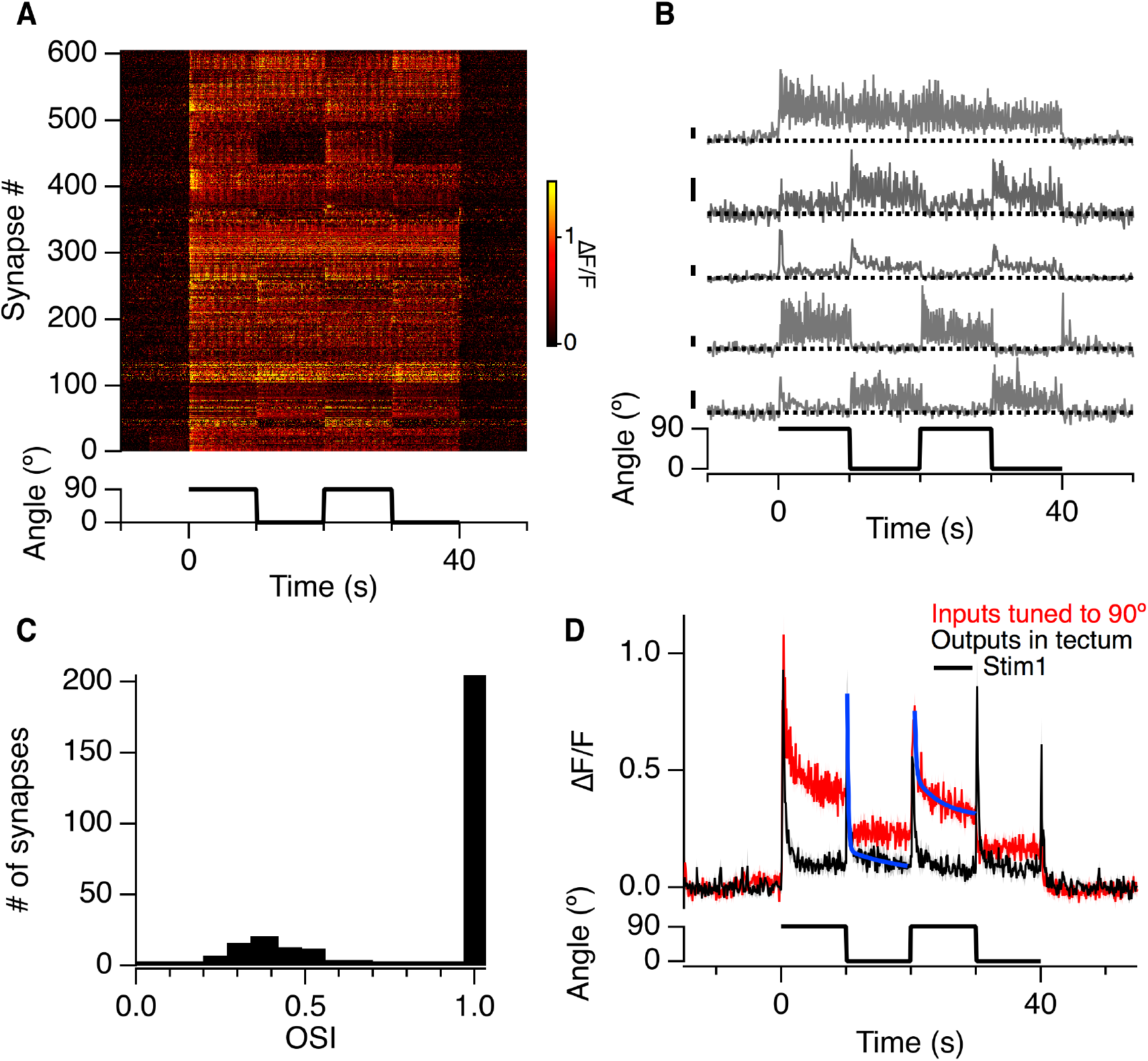
The output from bipolar cells displays orientation selectivity but not dynamic predictive coding. **A**) Raster plot showing the iGluSnFR signal of 606 bipolar cell inputs to 27 retinal ganglion cells in response to a full-field grating that switched orientation from 90° to 0° (stimulus as in Fig. 1). None of the synapses were activated at each change in orientation. **B**) Activity at five example synapses on RGC dendrites. The first is not sensitive to orientation while the lower four were stably tuned, preferring either the vertical or horizontal. All scale bars = 50% △F/F. **C**) Histogram of the OSIs for the orientation selective inputs measured in A. Note the two distinct populations. **D**) A comparison of the glutamatergic output of bipolar cell synapses tuned to the 90° (red; n = 151) and RGCs displaying dynamic predictive coding (black; n = 23). Shaded areas show ± sem. Note the differences in the time-course of adaptation. Solid blue lines are double exponential fits of the form y = y_o_ + A1((t − t_o_)/t_1_) + A_2_((t − t_o_)/ t_2_). The time constants for the bipolar synapses were 0.21 s and 4.2 s that accounted for 61% and 39% of the decay respectively. Whereas the tectal output was dominated by the fast component with a tau of 0.18 s accounting for 90% of the decay.

To identify RGCs sensitive to orientation we presented gratings that continuously reversed contrast at 5 Hz: keeping the temporal contrast fixed, the orientation of the grating was then switched from horizontal (90°) to vertical (0°) every 10 s until the grating was removed. In 102 of 106 RGCs the first presentation of the grating elicited a strong and transient output simply because of the increase in contrast (Fig. 1D). In 28 RGCs responses were insensitive to orientation (Fig. 1F), but in 66 the response to the grating depended on its orientation^31^. For instance, Fig. 1c shows a neuron generating a strong response to the vertical grating (R_0_) but very weak responses to the horizontal grating alone (R90). An orientation selectivity index (OSI) was defined as |R_0_-R_90_|/(|R_0_|+|R_90_|) and in the 66 of 106 RGCs classified as orientation selective this averaged 0.69 (Fig. 1E).

Of the orientation-selective RGCs, 43 exhibited relatively static tuning and adapted incompletely or not at all (Fig. 1G). But in the other 23 RGCs the orientation-selectivity was not fixed. Two examples are shown in Fig. 1H, where the neurons generated a transient glutamatergic output at both the horizontal-to-vertical and vertical-to-horizontal transitions. These outputs depressed by 92.0 ± 0.1 % within 0.26 ± 0.31 s, effectively suppressing the transmission of redundant information. Within ~9 s of this profound adaptation, these neurons had altered their tuning to allow the signaling of a future change in stimulus orientation, demonstrating the ability to generate a dynamic predictive code^9,12,13^. The retina of zebrafish therefore transmits information about orientation through two functionally distinct groups of RGC, either statically or dynamically tuned (Fig. 1G and H).

## Detection of spatial patterns originates in synapses of bipolar cells

To investigate how some RGCs dynamically alter their tuning to spatial patterns, we began by asking whether their excitatory inputs might be sensitive to orientation. Although electrical recordings in the soma of bipolar cells have not revealed any orientation selectivity^9^ it is known that inhibitory inputs onto the synaptic compartment can dramatically modify the electrical signal that drives transmission^22,39^. Imaging iGluSnFR provided the opportunity to directly assess the final output. The raster plot in Fig. 2A shows iGluSnFR responses from 606 synaptic inputs on the dendrites of 27 RGCs: 47% of these synapses displayed significant orientation selectivity and individual examples are shown in Fig. 2B. The distribution of OSIs measured for bipolar cell synapses displayed two distinct populations: the smallest group (30%) displayed a median OSI of 0.3, but the remaining 70% were almost perfectly selective for one orientation over the orthogonal (Fig. 2C). We confirmed this finding by surveying the whole population of bipolar cell synapses using zebrafish expressing a fast version of GCaMP6 fused to synaptopyhsin^40^: ~25% of these displayed orientation-selectivity when tested with either moving bars (Supplementary Figure 1) or the grating stimulus (Supplementary Figure 2) and there was a clear preference for vertically oriented stimuli (Fig. 3F). These results demonstrate that the analysis of orientation within the zebrafish visual system begins in the synaptic terminals of bipolar cells.

**Figure 3.**
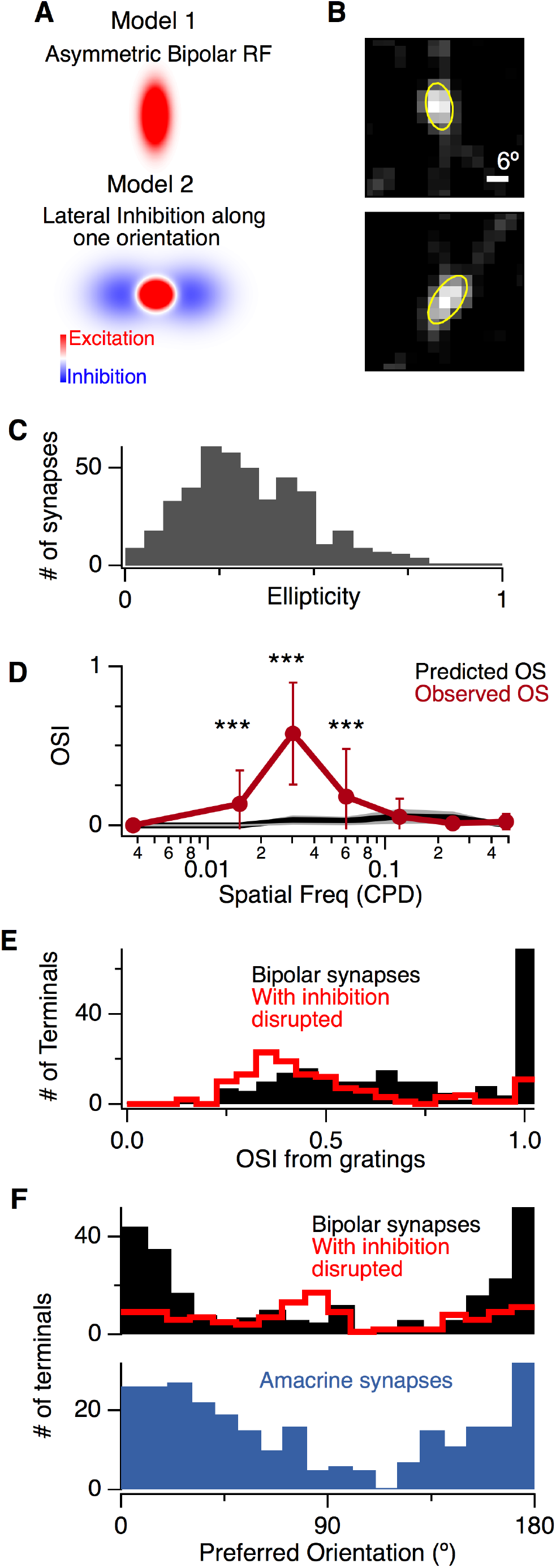
Intrinsic and extrinsic factors generating orientation selectivity in bipolar cell synapses. **A**) Two mechanisms potentially generating orientation selectivity. In Model 1 the synapse has an elongated receptive field center (red) which would lead to an orientation preference for vertical stimuli. Model 2 shows a bipolar cell receptive field (red) receiving inhibitory inputs that are stronger along the horizontal axis which would also lead to vertical selectivity in the bipolar cell synapse. **B**) Examples of receptive field centers mapped in synapses of two separate bipolar cells. Yellow lines show a fit with a 2D Gaussian used to estimate the length of their major and minor axes. **C**) Histogram of the receptive field ellipticity measured from 442 bipolar cell synapses. **D**) The OSI as a function of spatial frequency in bipolar cell synapses (red; n = 819) compared with the OSI predicted from the receptive field dimensions assuming linearity (black; n = 442). **E**) Comparison of the distributions of OSI under normal conditions (black, n = 257) and that remaining after disrupting inhibition with intravitreal injection of gabazine and strychnine (red, n = 127). **F**) *Top*: Distribution of preferred orientations under normal condition (black) and with inhibition disrupted (red). *Bottom*: Distribution of preferred orientations of 274 amacrine cell synapses showing significant OS. On average, amacrine cells were activated more strongly by the vertical (0°) compared to the horizontal.

The kinetics of the signals encoding orientation within the retina and the optic tectum were profoundly different. Fig. 2D compares the averaged output of RGCs displaying dynamic predictive coding (black) with the subset of bipolar cell synapses tuned to the horizontal (red). Bipolar cell outputs declined by an average of 49% over a 10 s period, in general agreement with the kinetics of contrast adaptation at the bipolar cell synapse measured using electrophysiology^41^ and the optical reporter sypHy^22,42^. In contrast, the output of RGCs adapted completely with a dominant time-constant of between 0.17 s and 0.25 s. Although depression intrinsic to bipolar cell synapses will also contribute to nulling the output to a constant signal^33^, this process is too slow and incomplete to account for the extent of adaptation in the signal delivered to the tectum (Fig. 2D). The rapid “zeroing” of the output in the face of an unchanging input is key to removing redundancy in a signal, and these results demonstrate that the retina carries out such an operation *after* the bipolar cell synapse.

## Intrinsic and extrinsic mechanisms generating orientation tuning in bipolar cell synapses

How do bipolar cell outputs become tuned to orientation? Two potential mechanisms, which might operate together, are shown in Fig. 3A. The first involves synapses with asymmetric receptive field centers causing features aligned with the longer axis to generate larger responses than those at other angles. We investigated this possibility by measuring calcium signals within bipolar cell terminals using SyGCaMP6f and mapping their receptive fields using the technique of filtered back projection^40^. Consistent with the dendritic field shapes of zebrafish bipolar cells^43^, the large majority of terminals had receptive fields displaying some degree of ellipticity (Fig. 3B). The median ellipticity was 0.30 (Fig. 3C), potentially providing a simple explanation for the population of synapses with OSI centered around 0.31 but not the second population with OSIs approaching one (Fig. 2C).

The second potential mechanism for generating pattern-detecting synapses invokes lateral inhibition from amacrine cells (Fig. 3A), which contact bipolar cell terminals through GABAergic connections^7^. This mechanism was first suggested when we measured OSIs using gratings of different spatial frequency and found that the receptive field centers of bipolar cell synapses were four-fold smaller than the spatial patterns that they signaled most strongly (Fig. 3D & Supplementary Fig. 2). The obvious candidates for neurons within the inner retina with large receptive fields are wide-field amacrine cells, so we tested their role by injecting a mixture of gabazine and strychnine into the intravitreal chamber to block both GABAergic and glycinergic inhibition. With inhibition blocked, only 10% of 1173 terminals displayed orientation selectivity, compared to 24.4% of 1053 terminals with inhibition intact (Fig. 3F & Supplementary Fig. 3). Notably, the population of bipolar cell terminals with OSIs around one was almost completely abolished (Fig. 3E), indicating that they were indeed generated by lateral inhibition. Blocking inhibition also flattened the distribution of preferred orientations from one in which terminals tuned to the vertical were strongly over represented to one in which horizontal orientations were represented at slightly higher frequencies (Fig. 3F).

Further evidence for the idea that inhibitory inputs contribute to the tuning of bipolar cell synapses is shown in Fig. 4, which plots SyGCaMP6f responses in terminals displaying differing degrees of selectivity for the vertical and horizontal. In the synapses with an OSI of one, a grating orthogonal to the preferred orientation often caused a significant *decrease* in calcium below the baseline i.e. a stimulus at the non-preferred orientation activated a counteracting inhibitory signal. These antagonistic responses depended on the spatial frequency of the stimulus (Supplementary Fig. 2) and were observed in 29% of terminals using a grating of 0.03 cycles per degree but only 9% of terminals using 0.121 cycles per degree, again indicating the involvement of amacrine cells with larger dendritic fields. When inhibition was blocked using gabazine and strychnine, responses with decreases in calcium for the null direction were completely abolished (not shown) and the distribution of preferred orientations was flattened (Fig. 3F).

**Figure 4.**
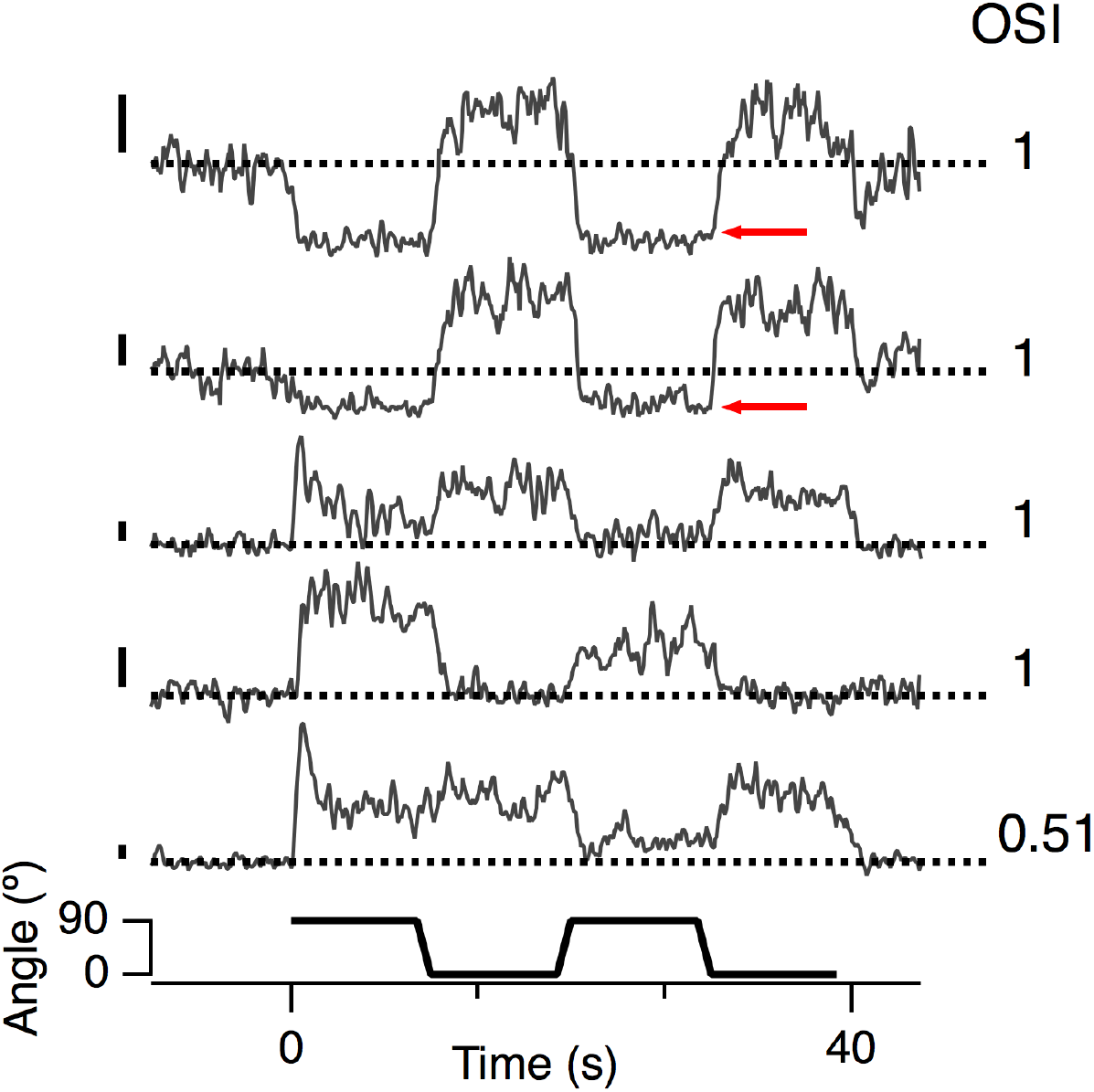
Lateral antagonism in the orientation domain. Examples of presynaptic calcium transients measured in bipolar cell synapses using SyGCaMP6f; same stimulus used in Figs. 1 and 2. The OSI for the top four terminals was one. The upper two terminals exhibited a fall in calcium reflecting hyperpolarization (red arrows) when the stimulus switched to the non-preferred orientation, indicating increased inhibition. Note that this measurement cannot distinguish increased inhibition from decreased excitation unless the inhibition is sufficient to decrease presynaptic calcium below basal levels. All scale bars = 50% △F/F

Direct evidence for the idea that amacrine cells can also be orientation-selective was obtained by measuring their synaptic activity using SyGCaMP3.5 under the *ptf1a* promoter^22^. Across a population of 982 synapses, 274 displayed a significant degree of orientation-selectivity with an over-representation of tuning to the vertical (Fig. 3F, lower panel). Amacrine cells are excited solely through bipolar cells, so one possibility is that they inherit their orientation-selectivity from these inputs. If this mechanism were common across the population of amacrine cells they would be expected to exhibit an average distribution of preferred orientations similar to bipolar cells and this was the observation (compare upper and lower panels in Fig. 3F). The distribution of preferred orientations in amacrine cells was, however, significantly broader than in bipolar cell synapses, likely reflecting the contribution of other factors such as intrinsic asymmetries in amacrine cell dendritic trees. Together, the results in Fig. 3 and 4 demonstrate that lateral antagonism in the orientation domain helps generate the most selective pattern detectors within the retina.

## The wiring of spatial pattern detectors to retinal ganglion cells

A key element of the pattern detector hypothesis is that individual RGCs receive inputs with a variety of tunings^9,32,33^. To investigate whether such RGCs exist in the retina of zebrafish, we used iGluSnFR to make a functional assessment of the rules governing bipolar cell to RGC connectivity. We sample 27 RGCs, measuring from an average of 22 synaptic inputs in each, distributed over several focal planes. Three general patterns of connectivity were observed. RGCs were either selective for untuned inputs (Fig. 5A; n = 6) or inputs tuned to the same preferred orientation (Fig. 5B; n = 6) or they received a mixture of excitation tuned to a variety of different orientations or none (n = 15). RGCs strongly selective for one of the two cardinal orientations (Fig. 5B) are likely to account for the population of RGCs with static tuning to orientation (Fig. 1G). Finally, of the RGCs receiving inputs of mixed properties, a subset of 3 displayed the basic wiring proposed by the pattern detector hypothesis - convergence of signals tuned to different orientations (Fig. 5C and D). Averaging across all the synapses sampled on each cell again demonstrated that these signals were slow to adapt and could not, therefore, account for the rapid suppression of the responses to a change in orientation measured at the RGC output (grey traces in Fig. 5C and D).

**Figure 5.**
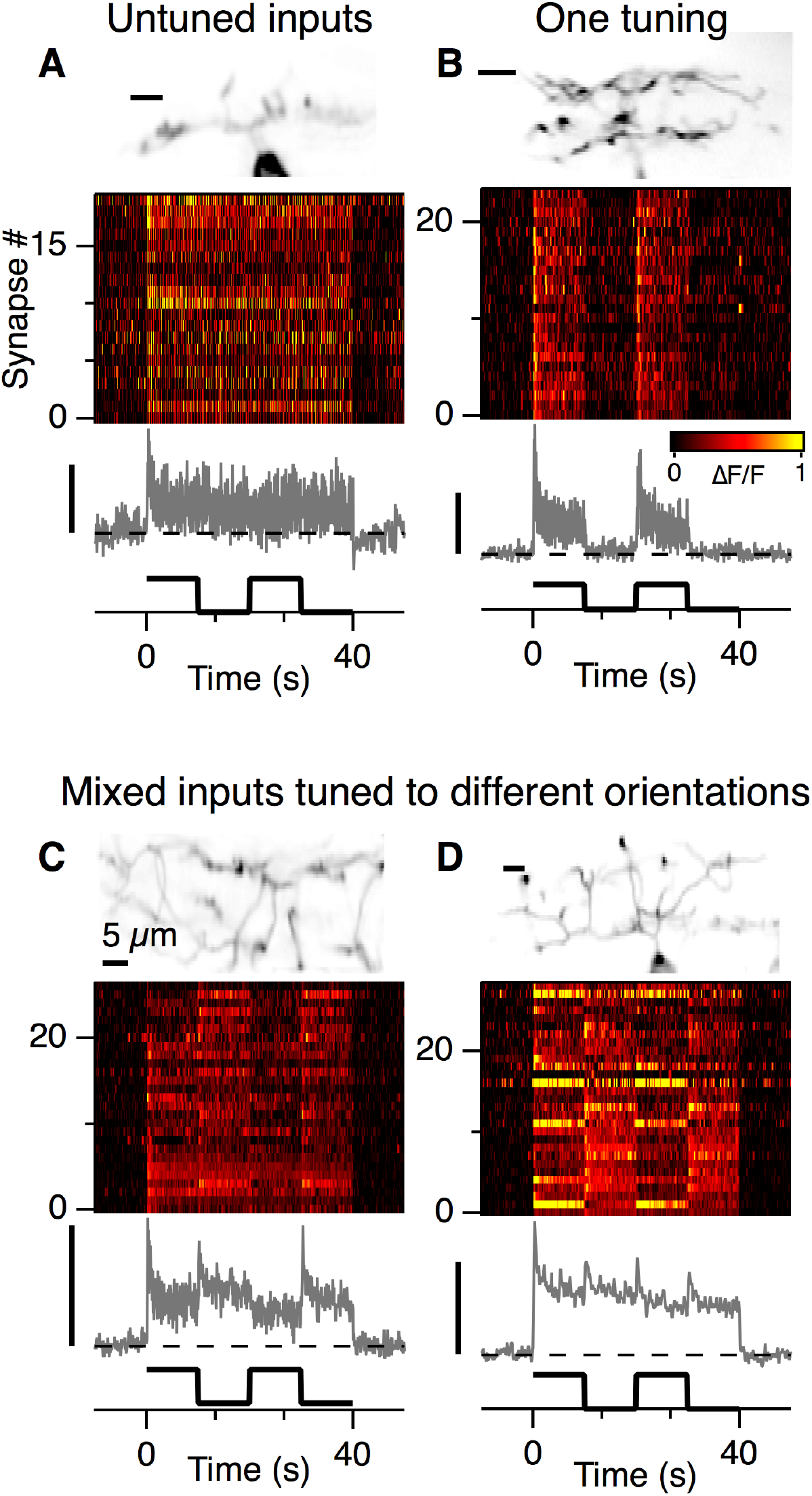
The wiring of bipolar cell synapses and retinal ganglion cells. **A**) An example of the synaptic inputs to one of 6 retinal RGCs that were selectively wired to non-OS synapses. The grey trace shows the average of these 19 inputs. **B**) An example of the synaptic inputs to one of 6 retinal RGCs that were selectively wired to synapses tuned to a similar orientation. Grey trace shows the average of these 24 inputs: the OSI averaged 0.98. **C**) Synaptic inputs to an RGC receiving a mixture 16 non-OS and 11 OS synapses. The net OSI averaged 0.55. **D**) An example of an RGC receiving a mix of excitatory inputs tuned to 0° (13), 90° (7) and non-OS (9). Note that the average excitatory drive to this cell shows a transient increase at each change in orientation. All image scale bars = 5μm and all △F/F scale bars = 50% △F/F

## A high-pass filter in retinal ganglion cells

To investigate further the transformation of the visual signal within individual RGCs we calculated the R^2^ value between iGluSnFR signals at the dendrites and axon, as would be used to assess a linear regression model. In 3 cells, the excitatory input alone was a very good predictor of the axons output accounting for ~66% of the variance, and two examples are shown in Fig. 6A. It appears that inhibitory signals received by these RGCs, either within the retina or tectum, play a relatively minor role in shaping the final output. But in the remaining five RGCs R^2^ averaged just 0.102 (Fig. 6B) and the output resembled a high-pass filtered version of the excitatory input: two examples are shown in Fig. 6C and E. These RGCs signaled changes in orientation strongly but attenuated the sustained DC component as well as the 5 Hz flicker, as shown by the spectrograms in Fig. 6D and F.

**Figure 6.**
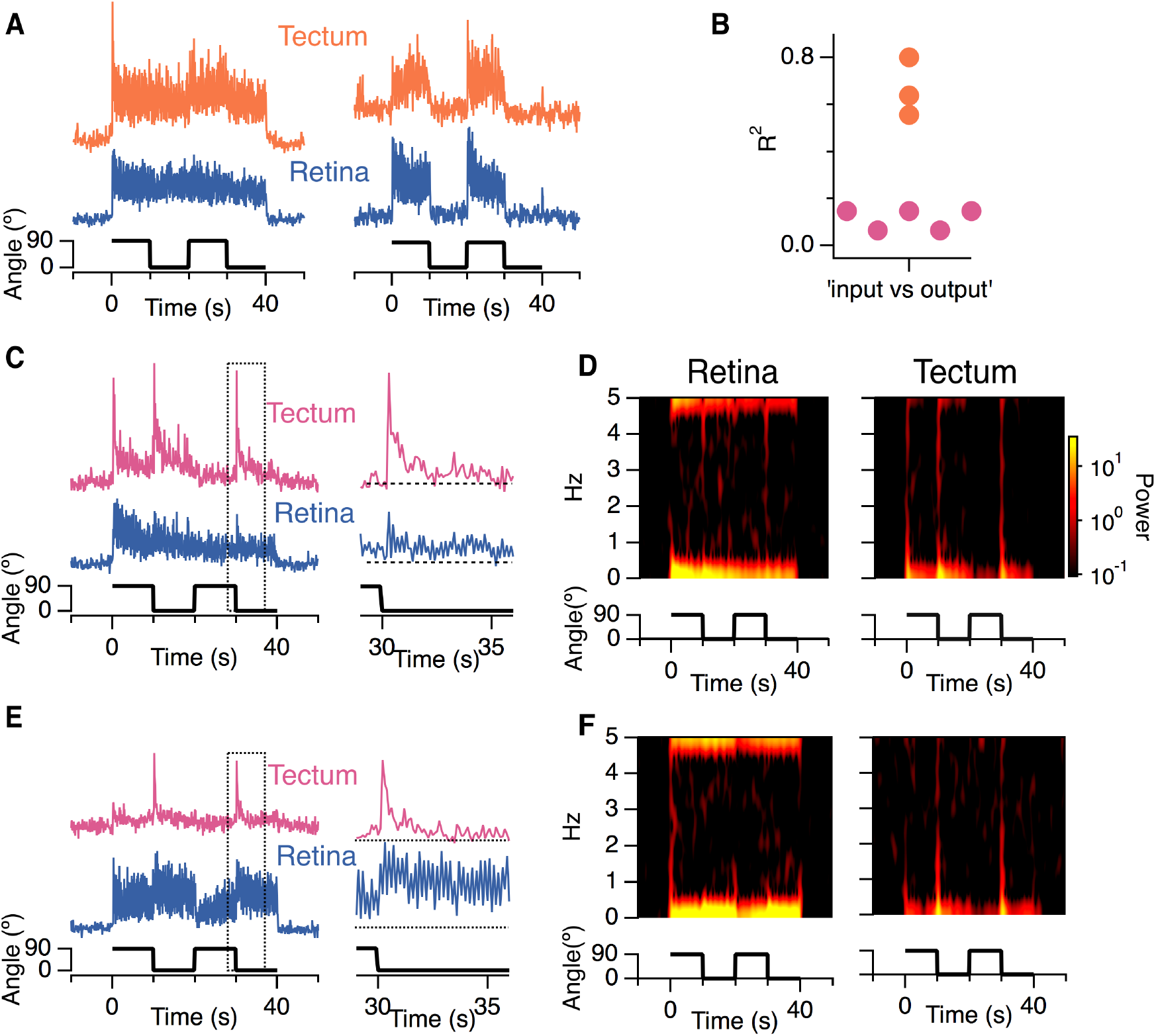
A high-pass filter operates in some RGCs. **A**) Examples of the averaged excitatory input (blue) and glutamatergic output (orange) of two RGCs in which input and output were strongly correlated. Stimulus as in Figs. 1 and 2. The RGC on the right was statically tuned (OSI = 1). **B**) R^2^ values for the correlation between the total excitatory drive and output for 8 different RGCs. The cells in A are marked in orange. The cells in C and E are marked in pink. **C & E**) Two examples of RGCs showing low degrees of correlation between excitatory input and output. The dashed box for each cell is blown up on the right, illustrating the filtering out of the sustained component of the response following a change in orientation. **D & F**) Spectrogram showing the power in the signals in C and E as a function of frequency (ordinate) and time (abcissa). Note that that the outputs in the tectum contain less power over a range of frequencies up to 5 Hz.

Might the high-pass filter operating in some RGCs reflect their connectivity to the other key element of the inner retina, the amacrine cells? A general feature of the retinal connectome is local feedforward inhibition (FFI), in which a bipolar cell excites both an RGC and an amacrine cell that inhibits the same RGC, often on the same dendrite^44–46^ (Fig. 7A). FFI is also a fundamental feature of the hippocampus and neocortex, where it controls the temporal window for firing in pyramidal neurons^47,48^ and can generate a high-pass filter^49^. Unfortunately, testing the role of FFI by pharmacological manipulation of inhibition was confounded by the simultaneous block of lateral inhibition causing large increases in the gain of excitation through bipolar cells. We therefore used a combination of electrophysiology and modelling to assess the role of FFI.

**Figure 7.**
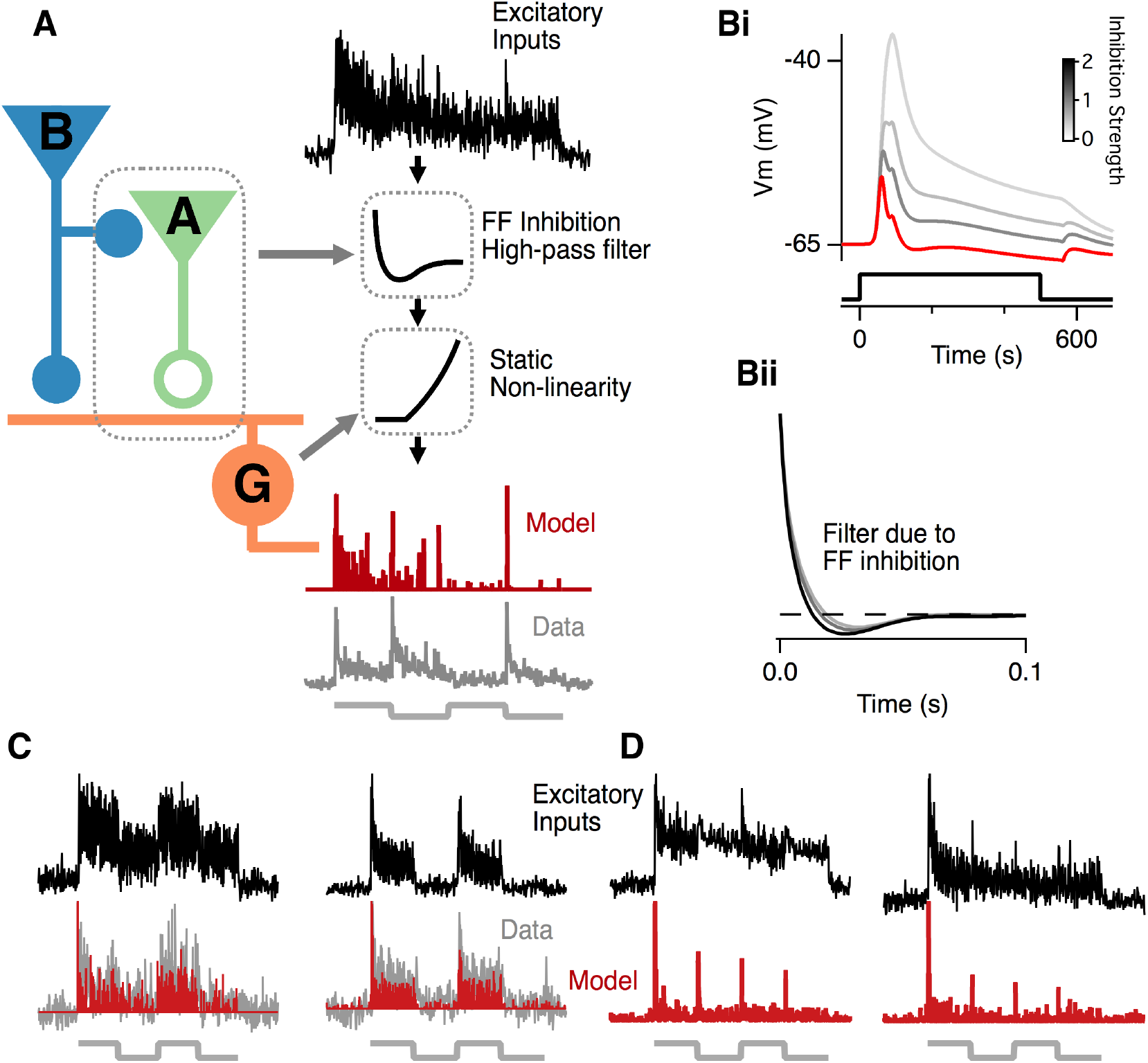
Feedforward inhibition can account for the transient outputs of RGCs generating a dynamic predictive code. **A**) A model in which an RGC (G) receives excitatory inputs from a bipolar cells (B) which also activates local feed-forward inhibition through an amacrine cell (A) contacting the same dendrite. The input to the model is the excitatory input measured using iGluSnFR (black trace) which is convolved with the high-pass filter calculated in B and then passed through a static non-linearity. The output of the model (red) can be compared with the output of the same RGC measured using iGluSnFR (grey). **B**) **i**. The membrane potential of an RGC in response to step activation of the excitatory and inhibitory synapses modelled in NEURON. Increasing the strength of inhibition makes the response more transient. **ii**) The impulse response of the temporal filter caused by feedforward inhibition shown in i. This filter assumed a 2:1 ratio of inhibitory to excitatory synapses (red trace in Bi). **C**) Two examples of the output of the model for RGCs receiving inputs tuned to a similar orientation. **D**) Two examples of the output of the model for RGCs receiving a mixture of inputs tuned to different orientations. The largest component of the signal is generated by changes in orientation.

A multicompartment model of a simplified RGC was made into which we integrated the properties of excitation and inhibition that we measured electrophysiologically^45^. Fig. 7Bi shows how excitation in the model RGC became briefer as the strength of the inhibitory input was increased. The ratio of inhibitory to excitatory synapses varies between RGCs, so we used the mean ratio of 2:1 measured experimentally in goldfish retina^45^ (Fig. 7Bii). The average excitatory input measured using iGluSnFR was then convolved with the high-pass filter introduced by FFI, with the results finally passing through a static non-linearity that represents thresholding performed by the spike generating mechanism in the RGC. A key assumption of this model was that the excitatory current injected into the RGC dendrite is directly proportional to the iGluSnFR signal, which has been confirmed experimentally^35^. The only parameter that was not empirically determined was the threshold for the non-linearity: in tests of the model shown in Fig. 7 we set this threshold to be 3 times the noise in the baseline. The results qualitatively reproduced the glutamatergic output measured in the tectum (Fig. 7A), including RGCs that were selectively wired to excitatory inputs of similar orientation selectivity (Fig. 7C) and those that received excitatory inputs tuned to different orientations and exhibited dynamic predictive coding (Fig.7D). These results demonstrate that the rapid and efficient suppression of an unchanging signal can be mediated by FFI.

## Discussion

A basic constraint on the design of neural circuits is the need to transmit information in an energy-efficient manner and removing redundancies from incoming signals is one of the most important ways to achieve this^2,4,50,51^. In this study we have delineated a circuit that allows individual neurons to signal changes in spatial patterns while strongly suppressing the transmission of unchanging and, therefore, redundant information. This implementation of a dynamic predictive code involves the re-tuning of orientation-selective RGCs^9^ by the circuit shown in Fig. 8, which contains the following basic elements: i) bipolar cell synapses acting as pattern detectors due both to their intrinsic orientation selectivity (Fig. 2 and Supplementary Figure 1) and to inhibitory inputs that act presynaptically to generate lateral antagonism in the orientation domain (Figs. 3 and 4); ii) a proportion of amacrine cells reflecting the orientation tuning of the bipolar cells that drive them (Fig. 3F); iii) individual RGCs receiving a mixture of excitatory inputs tuned to different orientations (Fig. 5D), and iv) feedforward inhibition through amacrine cells onto RGCs to generate a high-pass filter (Figs. 6 and 7).

**Figure 8.**
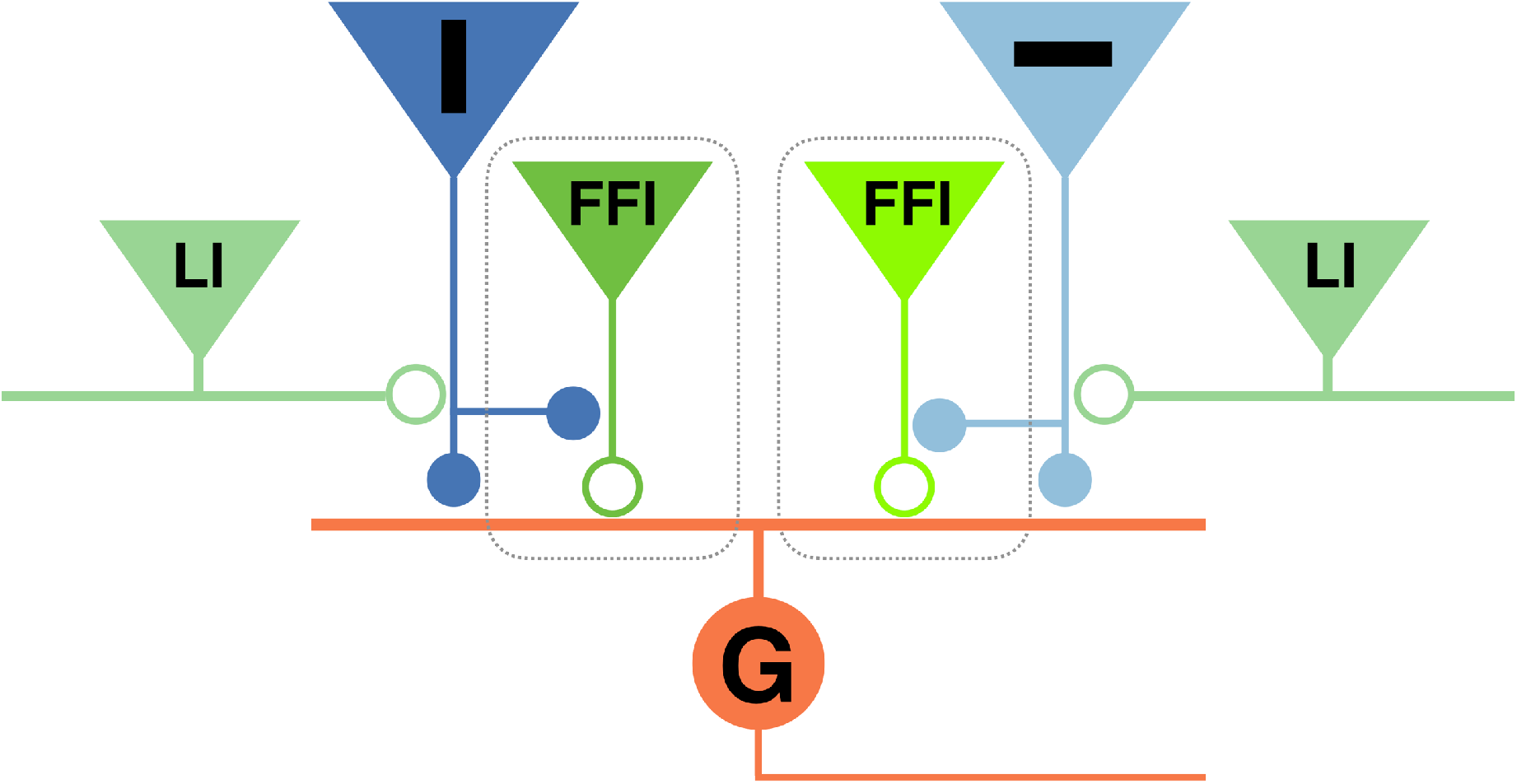
A retinal circuit generating a dynamic predictive code for orientated features. The RGC (G) receives a mixture of bipolar cell inputs tuned to different orientations, including those preferentially signalling the vertical (dark blue) and horizontal (light blue). Bipolar cell synapses tuned to orientation receive lateral inhibition from amacrine cells (LI) along their orthogonal orientation. Amacrine cells provide feedforward inhibition (FFI) onto the RGC dendrites generating a high-pass filter.

The basic features of the circuit in Fig. 8 are recapitulated in other parts of the brain that are thought to carry out some form of dynamic predictive coding, such as the primary visual^52,53^ and auditory^54^ cortices, the hippocampus^55^ and optic tectum^56^, and it may therefore reflect a general strategy by which excitatory and inhibitory neurons are connected to allow dynamic changes in the sensitivity of a post-synaptic neuron. For instance, individual pyramidal cells in V1 also receive excitatory synaptic inputs tuned to a variety of different orientations^30^ and their tuning shifts dynamically as they adapt to the distribution of orientations within a recent period of stimulation^24,26^. It is less clear whether changes in orientation tuning primarily reflect depression within excitatory synapses^24^ or the additional effects of inhibition^52^. In the retina, synaptic depression and FFI both reduce the net excitation caused by an input of a particular orientation, but with very different time-scales: presynaptic depression usually decreases the gain of an input over time-scales of several seconds (e.g Fig. 2D) while FFI acts on a fraction of a second^49^. Another key variable will be the strength of the feedforward inhibition received by the neuron acting as the pattern detector. We found a distinct subset of RGCs that did not act as high-pass filters (Fig. 6B), consistent with the idea that the FFI they receive is weak or non-existent. With variations, therefore, the basic circuit shown in Fig. 8 could act with varying efficiency and on different time-scales to alter tuning and suppress the transmission of maintained stimuli.

We found that most RGCs in zebrafish do not completely suppress an unchanging input, and do not, therefore, implement a dynamic predictive code in an ideal manner. Similarly, a survey in the retina of salamanders and rabbits found that only half of RGCs adapt to an orientated stimulus, with gain changes averaging a factor of about two^9^. The lack of more complete adaptation can now be understood in terms of the excitatory inputs that RGCs receive from bipolar cells, which depress incompletely and relatively slowly while a stimulus is maintained (Fig. 2). Modelling basic features of connectivity in the inner retina indicate that the steady excitatory input is only nulled effectively and rapidly in the subset of RGCs that additionally receive FFI (Figs. 6 and 7). The tectum therefore receives information about spatial patterns in at least two ways: some RGCs signal changes towards a preferred orientation and are stably tuned while others signal changes in any orientation and rapidly retune (Fig.1G and H).

It remains to be seen how these different signaling modes are used by downstream circuits. Answering such questions will be aided by measuring the distribution of spatial correlations in orientation space that a zebrafish is encountering in its normal environment of shallow, slow-moving, streams. Amongst the first pattern detectors – the bipolar cell synapses – there is an over-representation of vertical orientations (Fig. 1F) which might, for instance, reflect the importance of detecting vegetation growing upwards towards sunlight. RGCs generating a dynamic predictive code might not simply analyze changes in the external visual environment. In principle, they may also generate a signal that alerts the animal to rotations of its body axis relative to the surrounds, and in particular relative to the preferred (vertical) orientation. It is known that the righting reflex is driven by inputs from the visual system as well as the vestibular and somatosensory systems^57^ and signaling sudden changes in spatial patterns may serve to detect changes in body roll.

The integration of anatomical and physiological studies of neural circuits has been termed “functional connectomics”^58^, and this idea has generally been framed in terms of defining the rules that relate the functional properties of neurons with the synaptic connections they make. The present study highlights the importance complementing this information with an understanding of the dynamic properties of many individual synaptic connections. The use of fluorescent reporters of synaptic activity provides a promising approach for understanding one of the most basic aspects of any neural circuit, the input-output relations of the neurons within it.

## Online Methods

### Zebrafish

*Tg(-1.8ctbp2:SyGCaMP6)* and *Tg(ptf1a:gal4; UAS:SyGCaMP3)* fish were generated as described previously^1^. *Tg(10xUAS:iGluSnFR^ccu003t^*) was a kind gift by S. Renninger and M.Orger and was generated in the following way: DNA coding for iGluSnFR^2^, in the Gateway destination vector pDestTol2pA (http://tol2kit.genetics.utah.edu, ^3^), was cloned downstream of ten repeats of the Upstream Activation Sequence (10xUAS) using LR Gateway recombination. 12 ng/μl plasmid DNA and 40 ng/μl Tol2 transposase mRNA with 0.02% phenol red was injected into 1-cell stage embryos heterozygous for Isl2b:Gal4^4^. Transient and mosaic iGluSnFR-expression was achieved by injection of the HuCKalTA4 plasmid in combination with *Tol2* transposase^3^ into 1-cell stage embryos derived from in crosses of *Tg(10xUAS:iGluSnFR^ccu003t^*) which have been previously outcrossed to wildtype fish in order to split them from the Isl2b:Gal4 transgene. The concentration of the DNA and mRNA in the injection mix each were adjusted to 12.5 ng/ μl. Fish were raised and maintained under a 14 h light/10 h dark cycle and standard conditions as described previously^5^. To aid imaging through the eye, fish used for 2-photon imaging were either heterozygous or homozygous for the casper mutation, which results in hypopigmentation due to the lack of melano- and iridophores. To further suppress pigmentation fish were treated with 1-phenyl-2-thiourea (200 μM final concentration; Sigma) from 1 day post fertilisation to reduce pigmentation further. Fish were used at 6-8 days post fertilisation. All animal procedures were performed in accordance with UK Home Office guidelines and with the approval of the University of Sussex local ethical committee.

### Molecular Biology

The HuCKalTA4 plasmid has been generated via the Gibson Assembly Cloning method. The plasmid features the HuC promoter that drives expression of the optimized transcriptional activator KalTA4 in most neurons in the brain, including amacrine cells and RGCs in the zebrafish retina. The optimization of Gal4 has been described previously^6^. The plasmid contains a second small promotor (cmcl2) which drives expression of the mcherry fluorophore in the heart, this approach is commonly used in order to enable for phenotypic screening of transgenes that are not visible. The table shows the plasmid and primer information, the underlined nucleotide sequences show the part of the primer that anneals to the template during the polymerase chain reaction.

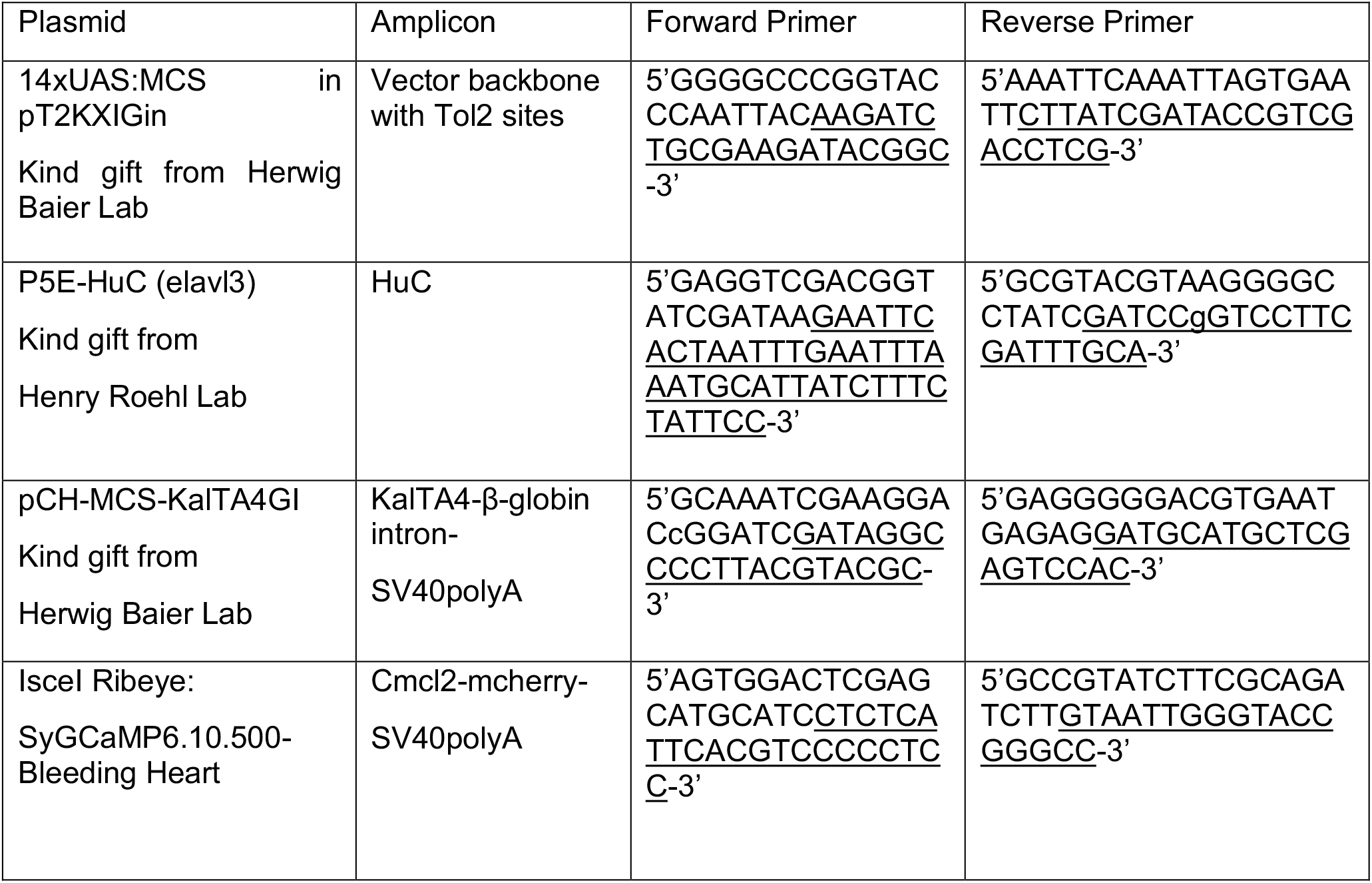

### Visual stimuli

Visual stimuli were generated using custom software written in Matlab (MathWorks, Natick, MA, U.S.A.) utilising Psychophysics Toolbox^7^. Stimuli were presented using an Optoma PK320 pico projector (Optoma, Watford, Hertfordshire, U.K.), modified so that only the red LED was used for projection^5^. The mean irradiance of the screen was 12.7 nW mm^−2^ and fish were positioned so that they viewed the centre of the screen and were adapted to the mean luminance for ≥10 mins. Square wave grating stimuli had an amplitude from −100% to +100% contrast and were designed so that the centre of a bar was always aligned with the centre of the screen, rather than an edge. For the spatial frequency tuning in Supplementary Fig. 2 and Fig. 3D each frequency was presented separately and in a pseudorandom order, the resulting videos were then concatenated in order of spatial frequency. For the moving bar experiments visual stimuli consisted of −100% contrast bars with a width 4.1° of visual angle which traversed the screen at 18.6° s^−1^ in 8 different directions, thus giving stimuli of 8 directions and 4 orientations. The bar height spanned the full size of the screen and stimuli were spaced 6 s apart. The sequence of 8 directions was presented in a pseudo random order then repeated 5 times. Visual stimulation was synchronised with imaging using custom-written code and a U3 LabJack digital-to-analog converter (Labjack, Lakewood, CO, U.S.A.).

### Two-photon imaging

Fish were immobilised in 3% low melting point agar (Biogene, Kimbolton, Cambs, U.K.) with one eye pointing at the middle of the screen. Bipolar and amacrine cell terminals and RGC dendrites in the central region of the retina were imaged at 10-20 Hz *in vivo* using a Scientifica 2-photon microscope (Scientifica, Uckfield, East Sussex, U.K.) equipped with a mode-locked Chameleon Ti-sapphire laser tuned to 915 nm (Coherent Inc., Santa Clara, CA, U.S.A.) with an Olympus XLUMPLANFL 20× water dipping objective (NA 1, Olympus, Tokyo, Japan). Emitted fluorescence was captured through the objective. Scanning and image acquisition were controlled under ScanImage v.3.8 software^8^. Locating the axon terminals of identified RGCs in fish with more than one labelled RGC was aided by the retinotopic distribution of RGC axons in the tectum ^9^ where dorso-temporal RGCs project to the ventro-anterior region of the tectum. We also confirmed that the axon terminal belonged to the identified RGC by laser ablating the RGC to check that the axon terminals stopped responding (Figure 1C), this was done by parking the beam over the cell body with the laser power at maximum, laser delivery was terminated as fluorescence dramatically increased, this coincided with destruction of the cell body.

## Analysis

### Image segmentation

Regions of interest (ROIs) corresponding to synapses were segmented from the registered time series using an iterative method, similar to ^10^. Initially we determined a local correlation map by cross correlating the time series of each pixel (x0) with its 8 neighbours (x1-8), with pixel x0 then being replaced by the maximum correlation value. This local correlation map was then used to seed the ROIs. The pixel with the highest value in the local correlation map provided the first ROI seed. This pixel had 8 nearest neighbours, each was tested for correlation with the seed and if a threshold was reached they were added to the ROI. This process was iterated for the neighbours of all pixels added to the ROI, when no further pixels were added the ROI was complete. The next ROI seed is chosen as the highest value from the remaining pixels in the local correlation map. Threshold values were chosen by the experimenter and were consistent across all fish analysed for a particular protocol. Only terminals that had responses >4*SD of the base line were included for subsequent analysis. For segmentation of iGluSNFR signals in the tectum: the same approach was used however care was taken to join ROIs whose time series was strongly correlated. This ensured that we did not attribute the output of one RGC giving rise to multiple synaptic terminals to outputs from multiple RGCs, but may mean that we underestimated the frequency of some responses types if separate cells generated two similar response types.

### Classification of retinal ganglion cells

Cells were classified as “dynamic predictive coding” if they responded with a transient on each transition that was greater than the steady state of the previous response by at least 4 standard deviations. RGCs were classified as orientation selective if the responses to the final two gratings differed by 4 standard deviations of their steady state responses. Note DPC and OS are not mutually exclusive classes.

### Analysis of orientation selectivity

Wherever stated the orientation selectivity index (OSI) was calculated using equation 1, where a and b are the responses to the last two orthogonal angles thus avoiding any contrast dependent adaptation (“responses” are defined below).

Equation 1:

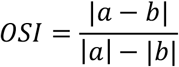

For iGluSNFR measurements for both the inputs and outputs of RGCs in response to gratings: Equation 1 was used to define OSI, again using the response to the latter two gratings. A response was only considered to have occurred for an orientation if the signal differed by 4 times the median absolute deviation (MAD) of the 15 s baseline. The value of a response was then determined in the following way: 1st we determined if the response was transient by comparing the maximum value measured within 0.5s after the transition, if this was >4*MAD of the median response over the whole 10s then the maximum value was used as the response, otherwise the response wasn’t considered transient and the median value was used. To prevent noise from generating spurious orientation selectivity, synapses were only classified as orientation selective if the responses to orthogonal orientations differed by greater than a multiple of the noise in the steady state response. This was 4*MAD for tectal responses and 3* MAD for bipolar synapses.

Equation 1 was used to define OSI for SyGCaMP measurements from bipolar cell and amacrine terminals in response to gratings. The amplitudes of the latter two grating where compared where “a” was the median amplitude during 10s at 90° and “b” was the median amplitude during 10s at 0°. Responses were only assigned an OSI if the difference between a and b was greater than twice the median-absolute-difference of the smaller response.

### Analysis of responses to moving bars

For SyGCaMP measurements from BPC and amacrine terminals we classified terminals as orientation-selective by computing a vector sum in orientation space for each of the 10 trials, terminals were then classified as OS based on the result of Moore’s version of Rayleigh’s test, with the critical value set to give a false positive rate of 1%. This critical value was determined by performing the same analysis on all terminals with the angle information shuffled; we then used the critical value that classified 1% of terminals as OS in this condition. Similar to previous reports in mouse, we did not find any significant direction preference in bipolar cell terminals^11^.

### Receptive field reconstruction

The receptive fields of bipolar cell terminals were mapped as described previously^5^, briefly a −100% contrast bar was flashed onto the retina with 3.2° spacing and at 5 angles ranging from 0° to 144°. Bars were presented in a pseudo random order and for 0.5 s with a 2 s duty cycle. These responses allowed the receptive fields to be accurately recovered with the filtered back projection method ^5^.

### Modelling the contribution of RF ellipticity to OSI

Phenomenological models were constructed in Igor Pro (Wavemetrics, Oregon, U.S.A.). 404 bipolar cell receptive fields were constructed as normalised 2D Gaussians with the major axis, minor axis and theta determined from fits to the 404 measured receptive fields (Figure. 3). Each model 2D Gaussian was first multiplied by a matrix representing a single grating stimulus, then the sum of all elements from the resulting matrix was rectified to give the response to a single grating. This was then repeated for the orthogonal orientation and equation 1 was used to calculate the OSI. This was repeated for all the spatial frequencies tested and the results shown in Figure 3D are the mean ±SD for all 404 receptive fields.

### Modelling the filter resulting from feedforward inhibition

We used the equations that fit previous measurements of the excitatory and inhibitory conductances from^12^ and placed a single excitatory and inhibitory synapse at the same point of a single dendrite of the simplified NEURON model^12^ used in Figure 7. The conductance of the excitatory synapse was set to 0.0005 nS and the conductance of the inhibitory synapse was varied from 0 to twice the magnitude of the excitatory synapse. To calculate the temporal filter resulting from the feed-forward inhibitory synapse we used normalised-least-mean-squares adaptive filtering with a filter length set to 750 ms. The filter was fit to 804 seconds of data generated by the simplified NEURON model which contained 1600 randomly timed synaptic events of varying duration.

### Drug application

To administer inhibitory antagonists to the retina in vivo, we injected ~4 nl of a solution containing 10 mM gabazine and 10 mM strychnine. We confirmed that these drugs gained access by including 1 mM Alexa 594 in the injection needle, within 5 mins of injection the dye could be detected within the inner plexiform layer of the retina (Supplementary Fig. 3A). However, the drugs do not distribute evenly within the eye, as can be seen in the accumulation of Alexa 594 in the intravitreal space.

## Supplementary Information

### 1. Orientation selectivity in bipolar cell synapses measured with moving bars

**Supplementary Figure 1.**
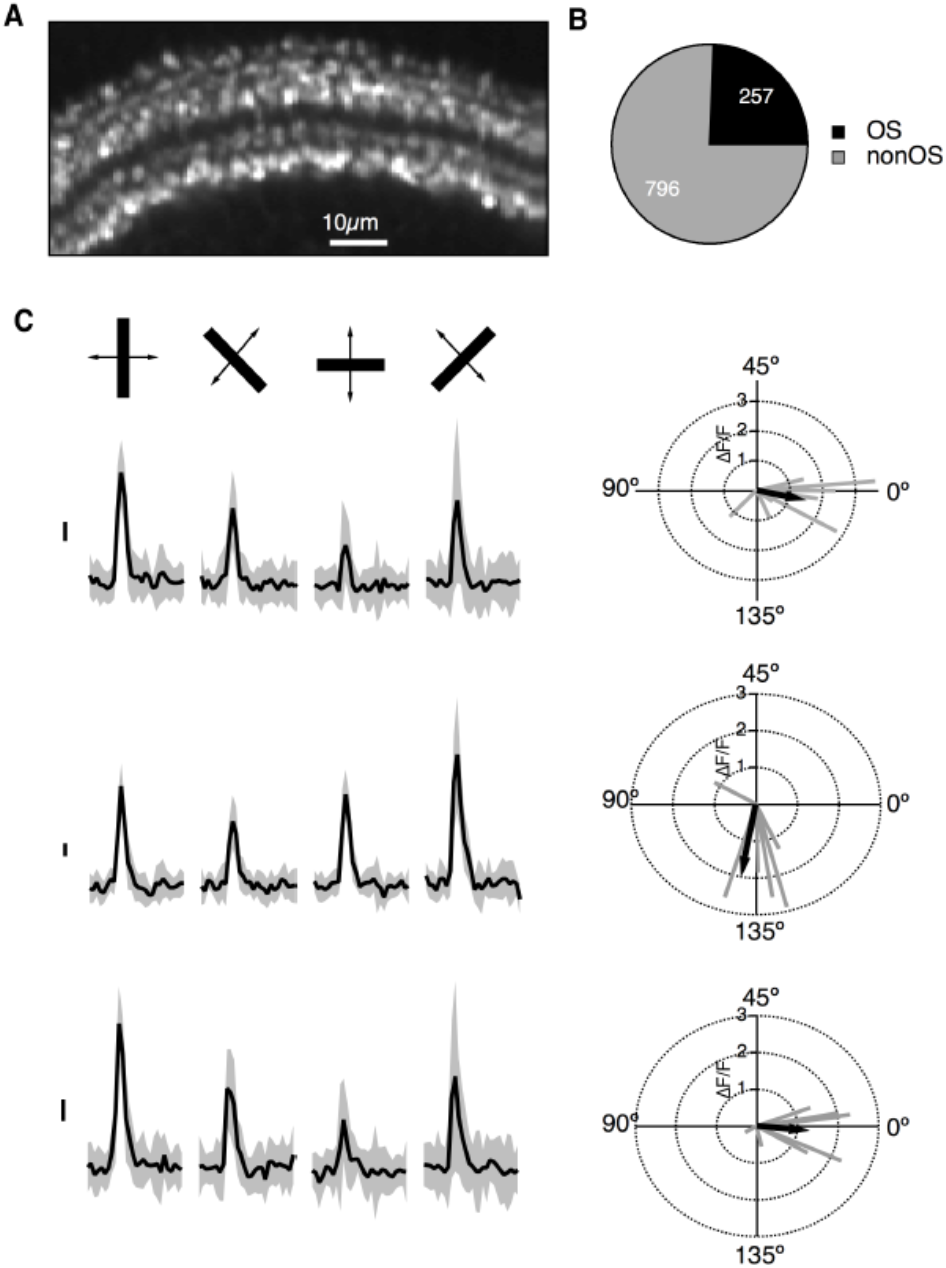
Orientation selectivity in bipolar cell synapses measured with moving bars. **A**) An array of bipolar cell terminals labelled with SyGCaMP6f imaged *in vivo*. **B**) 25% of 1053 measured synapses displayed a significant orientation preference with a false positive rate of 1%. The distribution of orientation preferences for these terminals are shown in Fig. 3F **C**) *Left* Example responses from 3 terminals to bars of different orientations moving across the field of view mean (black) ± 1SD (grey) of 10 repetitions. *Right*. The vector sum of each of the 10 trials plotted in orientation space (grey) with the average vector sum shown in black. To detect whether an individual terminal displayed a significant orientation preference we performed Moore’s version of Rayleigh’s test on the distribution of vector sums.

### 2. Spatial frequency tuning of bipolar cell synapses measured with SyGCaMP6f

**Supplementary Figure 2.**
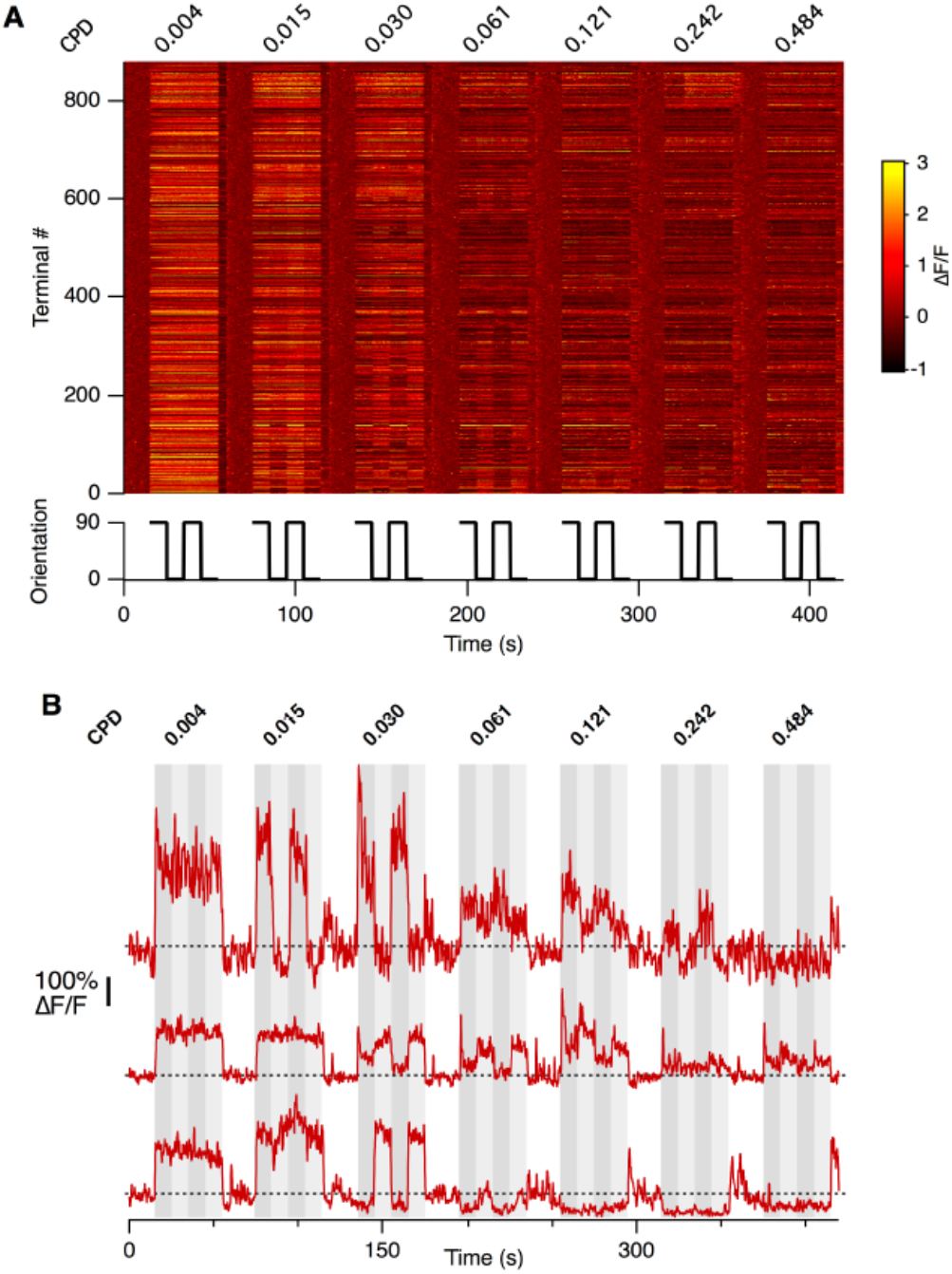
Spatial frequency tuning of bipolar cell synapses measured with SyGCaMP6f. **A**) The responses of 819 retinal bipolar cell synapses to gratings of spatial frequency ranging from 0.004 cycles per degree (CPD) to 0.484 CPD, 0.004 CPD is equivalent to full field. Each grating was given in a pseudo random order and then de-shuffled and concatenated for display purposes. This data generated the spatial frequency tuning shown in Fig. 3D. **B**) Examples from 3 bipolar terminals to gratings of varying spatial frequency.

### 3. Inhibition contributes to the orientation sensitivity of bipolar cell terminals

**Supplementary Figure 3.**
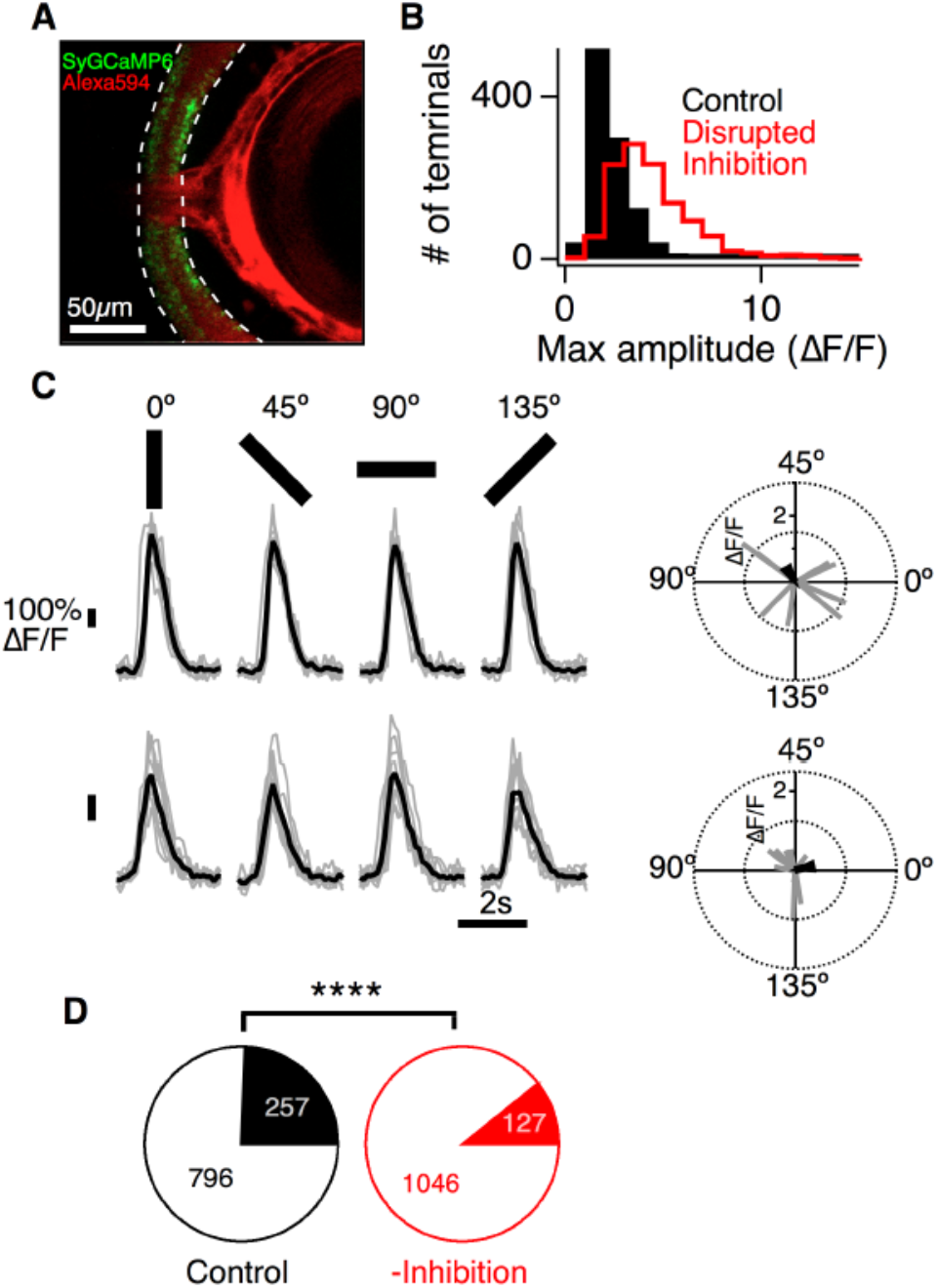
Inhibition contributes to the orientation sensitivity of bipolar cell terminals. **A**) Wide field of view of a zebrafish eye after an intravitreal injection of ~4 nl of a solution containing 10 mM strychnine, 10 mM gabazine and 1 mM Alexa 594. The inner plexiform layer is demarcated with dashed white lines with SyGCaMP6 expressing bipolar cell terminals labelled in green. The Alexa 594 signal is concentrated in the vitreous chamber and can be detected in the inner plexiform layer. **B**) Histograms comparing the average peak amplitudes of the response to the preferred stimuli for control (black, n=1053 terminals) and with intravitreal injection of inhibitory antagonists (red, n=1173, 7 fish). With inhibition blocked the responses of bipolar terminals tended to be larger (P<0.0001, Mann-Whitney test). **C**) Example responses from 2 terminals to bars of different orientations moving across the field of view, recorded in a retina with inhibitory antagonists blocked. Grey traces represent the 10 individual trials with the average shown in black. *Right*: The vector sum of each of the 10 trails plotted in orientation space (grey) with the average vector sum shown in black. **D**) The proportion of OS terminals was significantly lower in retinae where inhibition was blocked (red) when compared to control (black), P<0.0001, Fisher’s exact test).

## References

1 Barlow, H. B. in “Sensory Communication”, Possible principles underlying the transformations of sensory messages. (ed W. A. Rosenblith) 217–234 (MIT Press, 1961).

2 Srinivasan, M. V., Laughlin, S. B. & Dubs, A. Predictive coding: a fresh view of inhibition in the retina. Proceedings of the Royal Society of London. Series B, Biological sciences 216, 427–459 (1982).

3 Fairhall, A. L., Lewen, G. D., Bialek, W. & van Steveninck, R. R. D. Efficiency and ambiguity in an adaptive neural code. Nature 412, 787–792 (2001).

4 Sterling, P. & Laughlin, S. B. Principles of Neural Design. (MIT Press, 2015).

5 Atick, J. J. & Redlich, A. N. What does the retina know about natural scenes? Neural Comput. 4, 196–210 (1992).

6 van Hateren, J. H. Real and optimal neural images in early vision. Nature 360, 68–70 (1992).

7 Masland, Richard H. The Neuronal Organization of the Retina. Neuron 76, 266–280 (2012).

8 Kingston, A. & Autrusseau, F. Lossless image compression via predictive coding of discrete Radon projections. Signal Processing: Image Communication 23, 313–324 (2008).

9 Hosoya, T., Baccus, S. A. & Meister, M. Dynamic predictive coding by the retina. Nature 436, 71–77 (2005).

10 Nirenberg, S., Bomash, I., Pillow, J. W. & Victor, J. D. Heterogeneous Response Dynamics in Retinal Ganglion Cells: The Interplay of Predictive Coding and Adaptation. Journal of Neurophysiology 103, 3184–3194 (2010).

11 Ekman, M., Kok, P. & de Lange, F. P. Time-compressed preplay of anticipated events in human primary visual cortex. Nature Communications 8, 15276 (2017).

12 Sharpee, T. O. et al. Adaptive filtering enhances information transmission in visual cortex. Nature 439, 936–942 (2006).

13 Smith, E. C. & Lewicki, M. S. Efficient auditory coding. Nature 439, 978–982 (2006).

14 Sohoglu, E. & Chait, M. Detecting and representing predictable structure during auditory scene analysis. eLife 5, 19113 (2016).

15 Peyrache, A., Lacroix, M. M., Petersen, P. C. & Buzsaki, G. Internally organized mechanisms of the head direction sense. Nature Neuroscience 18, 569–575 (2015).

16 Parker, N. F. et al. Reward and choice encoding in terminals of midbrain dopamine neurons depends on striatal target. Nature Neuroscience 19, 845–854 (2016).

17 Schultz, W. Dopamine reward prediction-error signalling: a two-component response. Nature reviews. Neuroscience 17, 183–195 (2016).

18 Hindy, N. C., Ng, F. Y. & Turk-Browne, N. B. Linking pattern completion in the hippocampus to predictive coding in visual cortex. Nature Neuroscience 19, 665–667 (2016).

19 Stachenfeld, K. L., Botvinick, M. M. & Gershman, S. J. The hippocampus as a predictive map. Nature Neuroscience 20, 1643–1653 (2017).

20 Rao, R. P. & Ballard, D. H. Predictive coding in the visual cortex: a functional interpretation of some extra-classical receptive-field effects. Nature Neuroscience 2, 79–87 (1999).

21 Bastos, A. M. et al. Canonical microcircuits for predictive coding. Neuron 76, 695–711 (2012).

22 Nikolaev, A., Leung, K. M., Odermatt, B. & Lagnado, L. Synaptic mechanisms of adaptation and sensitization in the retina. Nature Neuroscience 16, 934–941 (2013).

23 Kastner, D. B. & Baccus, S. A. Insights from the retina into the diverse and general computations of adaptation, detection, and prediction. Current Opinion in Neurobiology 25, 63–69 (2014).

24 Muller, J. R., Metha, A. B., Krauskopf, J. & Lennie, P. Rapid adaptation in visual cortex to the structure of images. Science 285, 1405–1408 (1999).

25 Dragoi, V., Sharma, J. & Sur, M. Adaptation-induced plasticity of orientation tuning in adult visual cortex. Neuron 28, 287–298 (2000).

26 Benucci, A., Saleem, A. B. & Carandini, M. Adaptation maintains population homeostasis in primary visual cortex. Nature Neuroscience 16, 724–729 (2013).

27 Hubel, D. H. & Wiesel, T. N. Receptive fields of single neurones in the cat’s striate cortex. The Journal of Physiology 148, 574–591 (1959).

28 Ohki, K., Chung, S., Ch’ng, Y. H., Kara, P. & Reid, R. C. Functional imaging with cellular resolution reveals precise micro-architecture in visual cortex. Nature 433, 597–603 (2005).

29 Kondo, S. & Ohki, K. Laminar differences in the orientation selectivity of geniculate afferents in mouse primary visual cortex. Nature Neuroscience 19, 316–319 (2016).

30 Iacaruso, M. F., Gasler, I. T. & Hofer, S. B. Synaptic organization of visual space in primary visual cortex. Nature 547, 449–452 (2017).

31 Nikolaou, N. et al. Parametric functional maps of visual inputs to the tectum. Neuron 76, 317–324 (2012).

32 Lesica, N. A. et al. Adaptation to stimulus contrast and correlations during natural visual stimulation. Neuron 55, 479–491 (2007).

33 Gollisch, T. & Meister, M. Eye smarter than scientists believed: neural computations in circuits of the retina. Neuron 65, 150–164 (2010).

34 Graham, N. V. S. Visual pattern analyzers. (Oxford University Press, 2001).

35 Marvin, J. S. et al. An optimized fluorescent probe for visualizing glutamate neurotransmission. Nature Methods 10, 162–170 (2013).

36 Branco, T. & Hausser, M. Synaptic integration gradients in single cortical pyramidal cell dendrites. Neuron 69, 885–892 (2011).

37 Schmidt-Hieber, C. et al. Active dendritic integration as a mechanism for robust and precise grid cell firing. Nature Neuroscience 20, 1114–1121 (2017).

38 Borghuis, B. G., Marvin, J. S., Looger, L. L. & Demb, J. B. Two-photon imaging of nonlinear glutamate release dynamics at bipolar cell synapses in the mouse retina. The Journal of Neuroscience 33, 10972–10985 (2013).

39 Asari, H. & Meister, M. Divergence of visual channels in the inner retina. Nature Neuroscience 15, 1581–1589 (2012).

40 Johnston, J., Ding, H., Seibel, S. H., Esposti, F. & Lagnado, L. Rapid mapping of visual receptive fields by filtered back projection: application to multi-neuronal electrophysiology and imaging. The Journal of Physiology 592, 4839–4854 (2014).

41 Jarsky, T. et al. A synaptic mechanism for retinal adaptation to luminance and contrast. The Journal of Neuroscience 31, 11003–11015 (2011).

42 Odermatt, B., Nikolaev, A. & Lagnado, L. Encoding of luminance and contrast by linear and nonlinear synapses in the retina. Neuron 73, 758–773 (2012).

43 Li, Y. N., Tsujimura, T., Kawamura, S. & Dowling, J. E. Bipolar cell-photoreceptor connectivity in the zebrafish (Danio rerio) retina. J Comp Neurol 520, 3786–3802 (2012).

44 Chen, X., Hsueh, H. A., Greenberg, K. & Werblin, F. S. Three forms of spatial temporal feedforward inhibition are common to different ganglion cell types in rabbit retina. J Neurophysiol 103, 2618–2632 (2010).

45 Johnston, J. & Lagnado, L. General features of the retinal connectome determine the computation of motion anticipation. eLife 4, doi:10.7554/eLife.06250 (2015).

46 Diamond, J. S. Inhibitory Interneurons in the Retina: Types, Circuitry, and Function. Annu Rev Vis Sci 3, 1–24 (2017).

47 Pouille, F., Marin-Burgin, A., Adesnik, H., Atallah, B. V. & Scanziani, M. Input normalization by global feedforward inhibition expands cortical dynamic range. Nature Neuroscience 12, 1577–1585 (2009).

48 Isaacson, J. S. & Scanziani, M. How inhibition shapes cortical activity. Neuron 72, 231–243 (2011).

49 Milstein, A. D. et al. Inhibitory Gating of Input Comparison in the CA1 Microcircuit. Neuron 87, 1274–1289 (2015).

50 Barlow, H. B. in Sensory Communication (ed W. A. Rosenblith) 217–234 (MIT Press, 1961).

51 Niven, J. E. & Laughlin, S. B. Energy limitation as a selective pressure on the evolution of sensory systems. J Exp Biol 211, 1792–1804 (2008).

52 Priebe, N. J. Mechanisms of Orientation Selectivity in the Primary Visual Cortex. Annu Rev Vis Sci 2, 85–107 (2016).

53 Bock, D. D. et al. Network anatomy and in vivo physiology of visual cortical neurons. Nature 471, 177–182 (2011).

54 Li, L. Y. et al. A feedforward inhibitory circuit mediates lateral refinement of sensory representation in upper layer 2/3 of mouse primary auditory cortex. The Journal of Neuroscience 34, 13670–13683 (2014).

55 Mori, M., Gahwiler, B. H. & Gerber, U. Recruitment of an inhibitory hippocampal network after bursting in a single granule cell. Proceedings of the National Academy of Sciences of the United States of America 104, 7640–7645 (2007).

56 Luksch, H., Khanbabaie, R. & Wessel, R. Synaptic dynamics mediate sensitivity to motion independent of stimulus details. Nature Neuroscience 7, 380–388 (2004).

57 Kalueff, A. V. et al. Towards a comprehensive catalog of zebrafish behavior 1.0 and beyond. Zebrafish 10, 70–86 (2013).

58 Seung, H. S. Neuroscience: Towards functional connectomics. Nature 471, 170–172 (2011).

## References

1 Odermatt, B., Nikolaev, A. & Lagnado, L. Encoding of luminance and contrast by linear and nonlinear synapses in the retina. Neuron 73, 758–773, doi:10.1016/j.neuron.2011.12.023 (2012).

2 Marvin, J. S. et al. An optimized fluorescent probe for visualizing glutamate neurotransmission. Nature Methods 10, 162–170, doi:10.1038/nmeth.2333 (2013).

3 Kawakami, K. Tol2: a versatile gene transfer vector in vertebrates. Genome Biol 8 Suppl 1, S7, doi:10.1186/gb-2007–8-s1-s7 (2007).

4 Ben Fredj, N. et al. Synaptic activity and activity-dependent competition regulates axon arbor maturation, growth arrest, and territory in the retinotectal projection. The Journal of Nneuroscience 30, 10939–10951, doi:10.1523/JNEUROSCI.1556–10.2010 (2010).

5 Johnston, J., Ding, H., Seibel, S. H., Esposti, F. & Lagnado, L. Rapid mapping of visual receptive fields by filtered back projection: application to multi-neuronal electrophysiology and imaging. The Journal of Physiology 592, 4839–4854, doi:10.1113/jphysiol.2014.276642 (2014).

6 Distel, M., Wullimann, M. F. & Koster, R. W. Optimized Gal4 genetics for permanent gene expression mapping in zebrafish. Proceedings of the National Academy of Sciences of the United States of America 106, 13365–13370, doi:10.1073/pnas.0903060106 (2009).

7 Brainard, D. H. The Psychophysics Toolbox. Spat Vis 10, 433–436 (1997).

8 Pologruto, T. A., Sabatini, B. L. & Svoboda, K. ScanImage: flexible software for operating laser scanning microscopes. Biomed Eng Online 2, 13, doi:10.1186/1475–925X-2–13 (2003).

9 Robles, E., Laurell, E. & Baier, H. The retinal projectome reveals brain-area-specific visual representations generated by ganglion cell diversity. Curr Biol 24, 2085–2096, doi:10.1016/j.cub.2014.07.080 (2014).

10 Portugues, R., Feierstein, C. E., Engert, F. & Orger, M. B. Whole-brain activity maps reveal stereotyped, distributed networks for visuomotor behavior. Neuron 81, 1328–1343, doi:10.1016/j.neuron.2014.01.019 (2014).

11 Yonehara, K. et al. The first stage of cardinal direction selectivity is localized to the dendrites of retinal ganglion cells. Neuron 79, 1078–1085, doi:10.1016/j.neuron.2013.08.005 (2013).

12 Johnston, J. & Lagnado, L. General features of the retinal connectome determine the computation of motion anticipation. eLife 4, doi:10.7554/eLife.06250 (2015).

